# Major QTL controls adaptation to serpentine soils in *Mimulus guttatus*

**DOI:** 10.1101/328617

**Authors:** Jessica P. Selby, John H. Willis

## Abstract

Spatially varying selection is a critical driver of adaptive differentiation. Yet, there are few examples where the fitness effects of naturally segregating variants that contribute to local adaptation have been measured in the field. This project investigates the genetic basis of adaption to serpentine soils in *Mimulus guttatus*. Reciprocal transplant studies show that serpentine and non-serpentine populations of *M. guttatus* are genetically differentiated in their ability to survive on serpentine soils. We mapped serpentine tolerance by performing a bulk segregant analysis on F2 survivors from a field transplant study and identify a single QTL where individuals that are homozygous for the non-serpentine allele do not survive on serpentine soils. This same QTL controls serpentine tolerance in a second, geographically distant population. A common garden study where the two serpentine populations were grown on each other′s soil finds that one of the populations has significantly lower survival on this “foreign” serpentine soil compared to its home soil. So, while these two populations share a major QTL they either differ at other loci involved in serpentine adaptation or have different causal alleles at this QTL. This raises the possibility that serpentine populations may not be broadly tolerant to serpentine soils but may instead be locally adapted to their particular patch. Nevertheless, despite the myriad chemical and physical challenges that plants face in serpentine habitats, adaptation to these soils in *M. guttatus* has a simple genetic basis.

## INTRODUCTION

Natural landscapes are highly heterogeneous, resulting in selective pressures that differ between habitats. Such divergent selection can maintain genetic variation (Gillespie & Turelli, 1989; Levene, 1953), drive population differentiation (Felsenstein, 1976; Hedrick, 1986; Hedrick *et al.*, 1976) and ultimately promote speciation (Schluter & Conte, 2009). Some of the most striking examples of the power of natural selection to shape biological diversity involve adaptation of plants to extreme soil environments such as mine tailings, saline, acidic and serpentine soils (Brady *et al.*, 2005; Linhart & Grant, 1996; Mark R. Macnair, 1993; Rajakaruna, 2018). Evolutionary ecologists have studied plant adaptation to harsh soils for decades, providing some of the best examples of evolution in action (Linhart & Grant, 1996). Classic studies on plant adaptation to mine tailings demonstrated that plants can be locally adapted over a scale of meters despite substantial gene flow (Antonovics & Bradshaw, 1970; Jain & Bradshaw, 1966; McNeilly, 1968) and that local adaptation can lead to reproductive isolation (M.R. Macnair & Christie, 1983; McNeilly & Antonovics, 1968).

Transitions between soil habitats are often abrupt resulting in discrete habitat types with strongly divergent selective pressures. Large effect alleles may be more likely to be favored in such situations where populations are colonizing new habitats in which they are initially far from the fitness optimum (reviewed Dittmar *et al.*, 2016; Orr, 1998). Major genes have often been found for traits involved in adaptation to anthropogenic selection pressures such as industrial melanism (van’t Hof, 2011), warfarin resistance (Kohn, 2000) and insecticide resistance (Hemingway, 2004). These cases represent instances where selection is hard and individuals that lack appropriate genetic variation cannot survive. If individuals need a large phenotypic change to survive and reproduce this may favor the fixation of major genes (Mark R. Macnair, 1991). In addition, because distinct soil habitats often occur in close proximity if populations are locally adapted, this differentiation has likely evolved in the face of gene flow. Recent theoretical work shows that successful adaptation across discrete environments occurs either via few, large effect mutations or many small ones (Gilbert & Whitlock, 2017) and that selection for local adaptation with gene flow can lead to the clustering of multiple, linked loci (Yeaman & Otto, 2011; Yeaman & Whitlock, 2011). A number of recent QTL studies on metal tolerance and hyperaccumulation in plants growing in heavy metal contaminated soils suggests that adaptation to these stressful edaphic habitats may have a relatively simple genetic basis (e.g. cadmium and zinc tolerance in *Arabidopsis halleri* (Courbot *et al.*, 2007; Willems *et al.*, 2007) and *Thlaspi caerulescens* (Deniau et al., 2006); nickel tolerance in the serpentine endemic *Caulanthus amplexicaulis* (Burrell et al., 2012) and serpentine populations of *Silene vulgaris* (Bratteler et al., 2006); and copper tolerance in populations of *Mimulus guttatus* (K. M. Wright et al., 2013)). However, most of these QTL studies were conducted in lab-based conditions, often using hydroponic culture to isolate a single soil chemical variable thought to be important for adaptation. There are very few studies that have actually tested the fitness effects of loci that contribute to edaphic specialization in the field (but see Lexer *et al.*, 2003; K. M. Wright *et al.*, 2015) so we know little about the genetic architecture of adaptation to these habitats.

Serpentine soils are naturally occurring and present a diverse set of chemical and physical challenges to plants. These soils are widely distributed in western North America, stretching from the Baja peninsula to Alaska, but typically occurring in relatively small and isolated patches (Alexander *et al.*, 2007; Kruckeberg, 1984). While most terrestrial soils are derived from crustal rocks, serpentine soils are formed by the weathering of ultramafic rocks that originate in earth′s mantle (Alexander *et al.*, 2007). Because of this unique origin, serpentine soils are deficient in many essential plant nutrients [calcium (Ca), nitrogen (N), potassium (K), and phosphorus (P)] while also having elevated levels of magnesium (Mg) and heavy metals [nickel (Ni), cobalt (Co), chromium (Cr)]. In addition, these soils can be shallow and have lower water holding capacity than non-serpentine soils (Alexander *et al.*, 2007). This suite of factors makes serpentine soils inhospitable to many plant species resulting in sparse vegetative cover and habitats that are prone to erosion and elevated soil temperatures (Kruckeberg, 2002). The phrase “serpentine syndrome” was used by Jenny (1980) to emphasize the collective and often interacting effects of the chemical, physical, and biotic characteristics that make these soils such a difficult substrate for plant growth.

Despite this harsh and complex environment, serpentine habitats are home to unique plant communities with many endemic species. Serpentine endemism has long fascinated plant biologists; however, endemics may be reproductively isolated from their non-serpentine sister taxa, hindering genetic analysis of tolerance. Other species have populations that grow both on and off of serpentine soils and often show ecotypic differentiation (reviewed O’Dell & Rajakaruna, 2011). Field reciprocal transplant studies (Hufford *et al.*, 2008; Jurjavcic *et al.*, 2002; Kruckeberg, 1950, 1967; Sambatti & Rice, 2006; J. W. Wright *et al.*, 2006) and lab-based common garden experiments (Kay *et al.*, 2011; Kruckeberg, 1950; O’Dell & Claassen, 2008) typically demonstrate strong selection in serpentine soils with non-serpentine populations having higher mortality or greatly reduced growth relative to serpentine populations. Many studies testing for intraspecific differences use hydroponic culture to test the effects of isolated soil chemical features on plant growth. These experiments show that serpentine populations are primarily adapted to low Ca and a low Ca:Mg ratio (Brady *et al.*, 2005; Palm & Van Volkenburgh, 2014), but that in some species serpentine adaptation involves tolerance to high Mg (Proctor, 1970) or Ni (Burrell *et al.*, 2012; Gabbrielli *et al.*, 1990). However, caution is needed in interpreting hydroponic studies because these experiments may fail to replicate the complex interactions between different ions in the soil environment. Mg, Ca and Ni are all +2 cations and studies have revealed that differing concentrations of each affect plant availability of the other ions (Brooks, 1987; Gabbrielli & Pandolfini, 1984).

Despite the significant amount of work on serpentine adaptation, relatively little is known about the genetic basis of serpentine tolerance. The complexity of serpentine habitats suggests that changes at many loci might be necessary to adapt to these soils. Indeed, genome scans comparing serpentine and non-serpentine populations of *Arabidopsis lyrata* find dozens of highly differentiated SNPs (Turner et al. 2008, Turner et al. 2010, Arnold et al. 2016). In contrast, QTL mapping studies find that major genes contribute to elevated Ni tolerance in serpentine populations of *Silene vulgaris* and the serpentine endemic *Caulanthus amplexicaulis* var. *barbarae* (Bratteler et al. 2006, Burrell et al. 2012). Few QTLs of major effect were also shown to control reproductive versus vegetative investment which is thought to enhance drought escape on fast-drying serpentine soils in *Microseris douglasii* (Gailing et al 2004). These QTL studies focused on very specific traits differentiating serpentine and non-serpentine plants which is useful for elucidating important selective agents in these habitats but presumably presents a narrow view of the genetic basis of serpentine adaptation. QTL mapping approaches, in general, are limited in the genetic variants that are interrogated and the power to detect loci of small effect coupled with the overestimation of effects for detected loci (Beavis, 1994; Rockman, 2012). On the other hand, while genome scans can provide a more unbiased view of the loci under selection because these variants are not linked to relevant traits it is not known which of the outlier genes are most critical for tolerance and which may be subtle modifiers. Most importantly perhaps, none of the variants involved in serpentine adaptation identified by either approach have been tested for their fitness effects in the field so their true adaptive value is unknown.

The work presented here characterizes the genetic basis of adaptation to serpentine soils in *Mimulus guttatus* (Phrymaceae). *M. guttatus* is an outcrossing annual that grows throughout much of Western North America in seasonally wet soils. Across this range, *M. guttatus* displays tremendous ecological diversity and has become a model system for ecological genetic studies because of its tractability for experimental manipulation and a wealth of genetic and genomic resources (Hellsten *et al.*, 2013; Wu *et al.*, 2007). Populations of *M. guttatus* can be found in close proximity on and off serpentine soils across much of its range. No obvious morphological features distinguish populations growing on the different soil habitats and previous work has provided mixed evidence for genetic differentiation in edaphic tolerance between serpentine and non-serpentine populations. Two hydroponic studies grew populations from each soil type in low Ca:Mg conditions and did not find differential tolerance between serpentine and non-serpentine plants (Gardner & Macnair, 2000; Murren *et al.*, 2006). However, Palm *et al.* (2012) demonstrated that seedlings from a non-serpentine population do not survive past the juvenile stage when planted on native serpentine soil in the lab while a serpentine population has high survival.

Field-based reciprocal transplant studies are the best test for local adaptation. Here we present the results from three transplant studies carried out at different sites and in different years. Seedlings from multiple serpentine and non-serpentine populations as well as hybrid mapping populations (F2s or F3s) were planted at serpentine and non-serpentine field sites in California. Using multiple populations and conducting multiple experiments allows us to test for adaptation to the serpentine habitat, as opposed to highly local characteristics of a particular site or year. We use a bulk segregant approach to map a major QTL underlying survival differences at the serpentine field sites and show that this QTL has consistent effects across years and in two different populations. We follow-up this field work with a lab-based common-garden experiment in native serpentine soils to isolate the effects of selection due to soil variables and test how serpentine populations perform on foreign serpentine soils.

## MATERIALS AND METHODS

### Reciprocal transplant experiments

Seedlings from serpentine and non-serpentine *M. guttatus* populations were transplanted to serpentine and non-serpentine field sites to test for local adaptation. Three reciprocal transplant experiments were conducted: two at the Donald and Sylvia McLaughlin Natural Reserve (MCL) in 2010 and 2012 and a third at the Bureau of Land Management′s Red Hills Area (RH; Fig. 1) in 2010. Soil habitat of the seed collections used in these studies was established by a combination of field observations, USGS soil database designation for collection localities and, whenever possible, soil analysis (Fig. 1). Soil samples were taken by bulking soil from the top ∼8-12cm (roughly the rhizosphere of *M. guttatus)* from multiple locations throughout the population. These samples were analyzed for mineral nutrient content (see supplement for analysis details) and all serpentine populations had soil Ca:Mg levels below 0.35 while all non-serpentine populations were above 1 (sTable 1). Field seeds were planted in the greenhouse and crossed to derive full-sib families (1-2 families/population) for the 2010 experiments. The 2012 experiment used pooled seed created by combining equal numbers of seeds from 20 field collected maternal families/population. The MCL F2 mapping population was generated by reciprocally crossing the REM (serpentine) and SOD (non-serpentine) inbred lines and selfing the resultant F1s. In 2012 F3s from the same parental lines were produced by crossing 120 pairs of F2s and pooling ∼100 seeds from each cross. The RH mapping population consisted of outbred F2s generated by crossing two separate F1s that had unique inbred lines of the SLP (serpentine) and KFY (non-serpentine) populations as parents.

**Table 1.** Relative fitness and selection coefficients for each genotype at QTL on chromosome 13 on serpentine soil in four different experiments.

| **Experiment** | **Genotype** | **Survival**<br>**rate** | **Relative**<br>**Fitness** | **Selection**<br>**coefficient**<br>**(s)** |
| --- | --- | --- | --- | --- |
| **McL Field**<br>**2010** | SS | 25.1% | 1 | 0 |
|  | SN | 18.1% | 0.72 | 0.28 |
|  | NN | 0 | 0 | 1 |
| **McL Field**<br>**2012** | SS | 42.0% | 1 | 0 |
|  | SN | 34.1% | 0.81 | 0.19 |
|  | NN | 11.5% | 0.28 | 0.73 |
| **RH Field**<br>**2010** | SS | 22.3% | 1 | 0 |
|  | SN | 14.9% | 0.67 | 0.33 |
|  | NN | 2.5% | 0.11 | 0.89 |
| **RH Plates**<br>**Lab** | SS | 66.7% | 1 | 0 |
|  | SN | 42.7% | 0.64 | 0.36 |
|  | NN | 6.7% | 0.1 | 0.9 |

**Figure 1:**
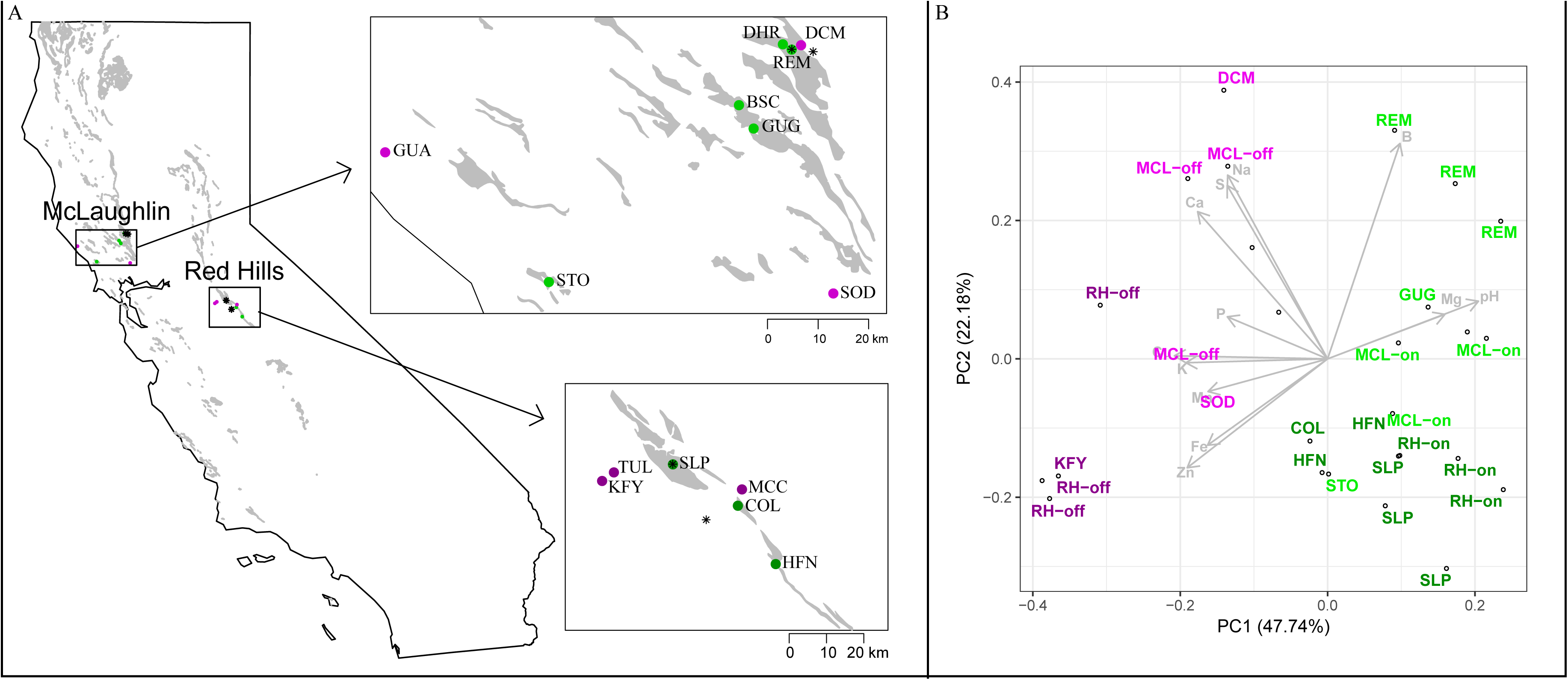
**A**) Locations of populations and field sites for reciprocal transplant studies. Serpentine localities are green while non-serpentine are pink. Red Hills localities are the darker shades. Transplant sites are marked with asterisks. Two serpentine transplant sites were used in each region but they are represented by a single point as, in both cases, sites are within 1km of each other. The REM and SLP populations are the home populations of one the serpentine transplant sites in McLaughlin and Red Hills respectively. Soil data obtained from *http://mrdata.usgs.gov/geology/state/state.php?state=CA* and serpentine soil distribution determined by extracting units with the rocktypes “peridotite” and “serpentinite” in R using packages rgdal, raster, rgeos and maps for shapefile manipulation and plotting. **B)** PCA biplot of soil variables for populations and field sites.

Three gardens were established for each experiment – 2 serpentine and 1 non-serpentine. However, dry conditions in 2012 resulted in heavy mortality at one of the serpentine sites prior to the first census; therefore, all details refer to the remaining two sites. Seeds were germinated on potting soil (Fafard 3B) outside at the McLaughlin Reserve in late January/early February and transplanted bare root to field plots 2 to 6 days after cotyledon emergence.

Transplants were randomized to cleared plots within native *M. guttatus* populations and marked with toothpicks. In 2010 small plots (4 × 3 seedlings, ∼ 7.5cm × 5 cm) were grouped together into blocks (12 plots/block with 6-8 blocks/site) and fifteen to thirty-five seedlings per family (sTable 1) along with ∼50 F1s and ∼500 F2s were transplanted to each field site. In 2012, seedlings were transplanted to cleared plots of 8×10 seedlings (∼35×45cm). Poor germination of some populations resulted in variable replication with 9-82 replicates/population/site (sTable 1). In addition, 800-1100 F3s were randomized within the 2012 plots. Planting date was recorded for all seedlings and survival time was calculated relative to planting date. In 2012, some late germinating individuals were transplanted to the field plots at the three-week census. The inclusion of these late transplants should not affect our survival analysis; if anything, they would make it more conservative.

Transplant survival as well as juvenile and adult size traits were recorded. Restrictions at both MCL and RH prohibited open pollination of the transplants so we were not able to collect more complete fitness data. In 2010 transplant survival was scored three weeks after planting and then weekly thereafter. In 2012 survival was scored at three and nine weeks after transplantation. Rosette diameter was measured 3, 4 and 5 weeks post-transplanting for the MCL2012, MCL2010 and RH2010 experiments respectively. In 2010, transplants were removed just prior to flowering, when the plants had buds with visible corolla tissue (for convenience we refer to this as “flowering date”). Height and length of the 1^st^ true leaf were measured at flowering. The 2010 experiments were terminated in early June when most plants had flowered or died (>90% at all sites). As most of the mortality on serpentine soils in the 2010 experiments and the lab-based common garden (see below) occurred shortly after transplanting, we concluded the 2012 experiment after 9 weeks in order to rescue a greater number of survivors for genotyping. None of the plants had flowered by this time so no measurements were taken. To collect tissue for genetic analyses, surviving F2/F3 individuals were shipped back to Duke, planted in potting soil and placed in the Duke greenhouses. Fresh bud tissue was collected from individual survivors as well as in bulk samples (taking one bud/F2) of serpentine and non-serpentine survivors for the MCL2010 experiment. We were not able to rescue all the F2s that survived to the end of the field experiments as some died after removal from the field plots but prior to tissue collection; however, the genotyped individuals appear to be representative of the overall group of survivors (see supplement).

Survival curves for serpentine, non-serpentine and hybrid plants in each planting habitat were constructed using Kaplan-Meier estimators calculated in with the package *survival* 2.38 (Therneau, 2015) in R version 3.3.3 (R Core Team, 2017). Survival time was calculated as days from transplantation to death. Plants that flowered and were removed from plots were treated as censored data as were plants still alive at the end of the experiment. We used log-rank tests to analyze cumulative differences in the survival functions between all classes (serpentine, non-serpentine, F1 and F2/F3) in each habitat. Significant overall log-rank tests were followed by post-hoc pairwise comparisons to see which groups were significantly different. We also looked at survival differences by population as well as tested for cytoplasmic effects on survival in the F1s and F2s.

Differences in plant size and days to flowering between plants from serpentine and non-serpentine populations were analyzed using analysis of variance (ANOVA). We first checked trait correlations and leaf length and height were highly correlated (MCL2010 r = 0.81; RH2010 r= 0.75) so we only include height in the subsequent analyses. While rosette diameter and height were also highly correlated (MCL2010 r=0.50; RH2010 r = 0.60) we analyze both traits separately as rosette diameter was measured early in the season and therefore scored on more plants than height. Both rosette diameter and height are negatively correlated with days to flower (MCL2010 r = −0.35, −0.33; RH2010 r = −0.31, −0.11 for rosette and height respectively). Boxplots of rosette diameter, height and days to flower for all plant classes in both habitats are provided in the supplement. However, high mortality at the serpentine sites resulted in unbalanced design so we restrict our formal analyses to the non-serpentine sites where we used two-way ANOVA to test for the main effect of habitat of origin while controlling for block effects. Tests were carried out in R using the lmer() function from the *lme4* package (Bates *et al.*, 2015) and block was treated as a random factor. Type II Wald F tests with Kenward-Roger degrees of freedom calculated with the Anova() function from the package *car (Fox & Weisberg, 2011)*. Rosette diameter and height were log-transformed to satisfy residual normality assumptions.

### QTL mapping

To rapidly map QTLs controlling survival differences on serpentine soil we performed a bulk segregant analysis (BSA; Michelmore *et al.*, 1991) with the F2 survivors from the MCL2010 experiment. Bulk DNA samples collected from the serpentine and non-serpentine survivors were sequenced on the Illumina platform to generate allele counts at SNPs across the genome. DNA was extracted using a urea protocol modified from Shure *et al.* (1983), submitted to the Duke Genome Sequencing and Analysis Core Resource for library preparation, and sequenced on the Illumina GAII for 75bp SE reads. To improve coverage in the serpentine survivor pool, the DNA was later re-submitted for library preparation and sequencing on the Illumina HiSeq2000 for 100bp SE reads. The inbred parental lines (REM and SOD) were sequenced for 150PE reads on the HiSeq2000 following library preparation with the Nextera DNA Library Prep Kit (Illumina, San Diego, CA, USA) from DNA extracted with the GeneJET Genomic DNA Purification Kit (Thermo Fisher Scientific, Waltham, MA, USA).

Raw reads were checked for quality using FastQC (Andrews, 2010) and then trimmed for quality and adapter sequence with TrimGalore (version 0.4.0 with cutadapt version 1.8.3; Martin, 2011) on default settings except for stringency --5. The quality scores for the reads from the Illumina GAII platform were adjusted to Sanger encoding using seqtk (version 1.2; Li, 2012). Trimmed reads were then mapped to the *M. guttatus* reference genome v2.0 (Hellsten *et al.*, 2013) using the BWA (version 7.15) mem algorithm with default settings (Li & Durbin, 2010).

Bam file cleaning and duplicate marking was performed using PicardTools according to GATK′s best practices (McKenna *et al.*, 2010; Van der Auwera *et al.*, 2002). Variants were called using GATK′s HaplotypeCaller (version 3.4) followed by joint genotyping on all four samples – two F2 bulks and two parental samples – using GenotypeGVCFs. Indels and repetitive regions (identified using a bed file created from the hardmasked genome available on phytozome) were removed from the variant file using vcftools (version 0.1.14; Danecek *et al.*, 2011). Finally, SNPs were thinned in vcftools to remove variants that were within 100bp of each other.

Bulk segregant is based on the expectation that alleles frequencies in the two pools will be similar across the genome but diverge in regions that contain QTLs. However, comparing allele frequencies does not account for the overall differences in depth of coverage between our two bulks nor the random variation in sequencing depth. Thus, we use the G-statistic based on allele counts in each pool to quantify the differentiation between our bulks as it accounts for the weight of evidence related to sample sequence depth (Magwene *et al.*, 2011). To generate a list of high-confidence SNP markers we filtered the raw variant calls for mapping quality ≥ 30 and depth ≥ 3 and ≤ 95^th^ percentile (DP ≤ 18 non-serpentine pool; DP ≤ 48 serpentine pool). Both parental lines had raw coverage around 30x and were filtered to retain sites with 12 ≤ DP ≤ 90. We restricted our analysis to sites where the F2 pools were segregating and the parental lines were fixed for alternate alleles. The allele calls in the parental samples were used to polarize SNP alleles in the F2 pools. Because of low coverage in the F2 pools (mean coverage post-filtering non-serpentine = 7.5x; serpentine pool = 18x), we summed counts of serpentine and non-serpentine alleles in each pool across windows of 25 SNPs. Windows larger than 1Mb were excluded and G was calculated from the windowed allele counts.

Putative QTLs identified via BSA were confirmed by genotyping individual F2s at PCR-based markers and comparing patterns of segregation distortion between the serpentine and non-serpentine survivors. DNA was extracted from the F2s and parents using a modified CTAB protocol (Kelly & Willis, 1998). The parents were screened for exon-primed intron-crossing markers derived from expressed sequence tags (Fishman *et al.*, 2008). Polymorphism was evaluated in terms of length variation of the PCR products, typically caused by indel variation in the introns. Amplified products were run on an ABI 3730xl DNA Analyzer (Applied Biosystems, Foster City, CA, USA) for fragment analysis and fragment size was scored using

Genemarker (SoftGenetics, State College, PA, USA). All F2 survivors from the MCL2010 experiment were genotyped at several markers in putative QTL regions (Fig. 3B and sTable 4). Markers were tested for goodness of fit with Mendelian (1:2:1) expectations using chi-square tests. The F2/F3 survivors from the RH2010 and MCL2012 experiments were also genotyped at markers within QTL identified in the MCL2010 mapping population.

### Lab-based common garden experiment

To test whether field survival differences were due to soil properties as opposed to other characteristics of the serpentine habitat (e.g. water availability, exposure, community composition) we conducted a common garden experiment growing serpentine and non-serpentine plants on serpentine soil in lab conditions. Inbred lines from two on/off pairs (REM/SOD and SLP/TUL; Fig. 1) as well as F1s and F2s for each pair (except for the SLPxTUL F1 which had limited seeds) were grown on serpentine soil collected from each of the serpentine localities. The RH soil consisted of combined samples collected from each of the plots in the field experiment above. The MCL soil was collected ∼0.8km away from the home site of the serpentine parent. The same soil was used by Palm *et al.* (2012) and they show that it is not significantly different in chemical composition from the soil at the parental site. The soils were sifted through a 2mm mesh screen and sterilized (121°C, 15psi, 90min).

Seeds were planted on 60×15mm petri plates, covered with ultrapure water (Nanopure Diamond purified) and stratified in the dark at 4°C for 5 days. Plates were then moved to a growth chamber (12 hours light; day/night temperatures of 22°C/18°C) and after radicle emergence (∼2days) transferred with a cut-tip pipette to petri plates filled with serpentine soil. Twelve seeds were planted per plate with 2-8 replicate plates/soil type for the parental lines and F1s and 40 plates per soil type for each of the F2s (20 plates per direction of the cross). Seedlings were checked daily and watered with ultrapure H_2_O when the soil surface was dry. Survival was scored weekly for five weeks. Final survival differences between lines were tested with G-tests of independence on overall counts of “dead” versus “alive.”

## RESULTS

### Survival differences at serpentine field sites

There were significant differences in survival between serpentine and non-serpentine plants at the serpentine field sites in all three reciprocal transplant experiments (Fig. 2). Very few non-serpentine plants survived to flower at the serpentine sites - MCL2010 1/91; RH2010 4/253; MCL2012 2/91 – while plants from serpentine populations enjoyed intermediate survival rates. The survival rate for the F1s in both 2010 experiments was not significantly different from that of the serpentine populations indicating that serpentine tolerance is dominant. The F2s/F3s had intermediate survival rates that were significantly different from both the serpentine and non-serpentine classes in 2010 but in 2012 pairwise log-rank tests show no overall difference between the F3s and the serpentine plants. There were no significant survival differences at the non-serpentine sites except for the MCL2012 experiment where the F3s had higher survival than both the serpentine and non-serpentine plants.

**Figure 2:**
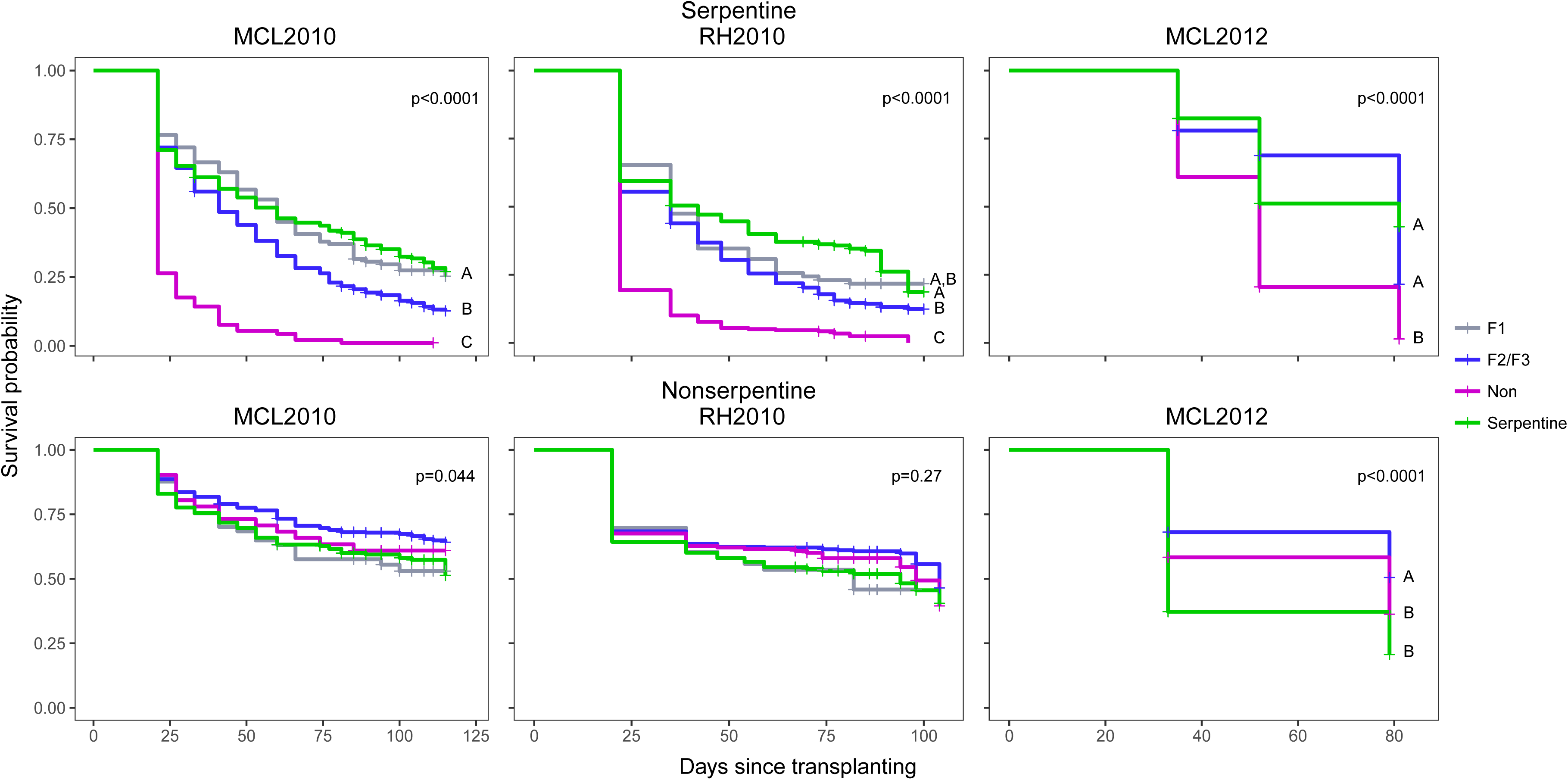
Kaplan-meier survival curves for serpentine, non-serpentine, F1 and F2/F3 plants in serpentine (top row) and non-serpentine (bottom row) field sites. P-values given for log-rank tests of all groups. For experiments with significant overall survival differences pairwise post-hoc log-rank tests were conducted and groups connected by the same letter are not significantly different. MCL2010 nonserpentine none of the pairwise posthoc tests were significant after Bonferroni correction.

Principal component analysis of soil chemical variables for field sites and population home sites separates serpentine and non-serpentine localities along PC1 and regions (McLaughlin versus Red Hills) along PC2. PC1 explains 48% of the overall variation and all the variables except for boron (B) have similar loadings (sTable 2) emphasizing the complex nature of the differences between these habitats. Not surprisingly the serpentine soils have higher levels of Mg and lower levels of nearly all other elements analyzed (Fig. 1). The regional differences along PC2 indicate that McLaughlin soils have higher levels of B, Ca, sodium (Na) and sulfur (S) while Red Hills soils have higher levels of zinc (Zn) and iron (Fe). However, there is a fair bit of variation within these regional habitat clusters and even among samples from the same sites (e.g. REM which is both a population and one of the MCL serpentine transplant sites). This soil variation between localities may explain some of the survival differences between serpentine populations at the serpentine field sites (Fig. S1). While all the serpentine populations enjoyed significantly higher survival than the non-serpentine populations, some serpentine populations had higher survivorship rates than others: for example, the home RH serpentine population (SLP) had the highest survival of the three serpentine populations included in the RH2010 experiment. These differences could be due to local variation in soil chemistry or to other factors, such as soil physical properties, water availability or exposure.

### Size differences at non-serpentine sites

While there were no survival differences at the non-serpentine sites there were differences in plant size, wherein plants from non-serpentine populations were larger than those from serpentine populations (Fig. S2) indicating a potential cost to tolerance. In 2010 there were significant differences in rosette diameter between serpentine and non-serpentine plants at the non-serpentine field sites (RH F_1,178_=6.71, p=0.01; MCL F_1,173_ = 4.245, p=0.041). However, there we did not detect differences in rosette diameter in 2012 (F_1,71_=0.621, p = 0.433) possibly due to the lower sample size and earlier measurement date. Height and flowering time were only measured for the 2010 experiments. There were significant height differences between serpentine and non-serpentine plants in the RH experiment (F_1,139_=4.73, p =0.031) but not at MCL (F_1,99_=0.05, p=0.824) and there were no significant differences in flowering time in either experiment (MCL F_1,103_=0.258, p = 0.612; RH F_1,141_=0.069, p= 0.793). An overall effect of planting habitat is evident in Fig. S2 where plants at the serpentine sites are smaller and flower slightly earlier.

### Major QTL contributes to serpentine survival

Using a bulk segregant approach we identified a single region of the genome that contributes to survival on serpentine soils. Sequencing pools of the F2 survivors from the MCL2010 experiment identified a QTL on the end of chromosome 13 displaying an elevated G-statistic relative to the rest of the genome (Fig. 3A). In this region, the serpentine allele frequency in the serpentine survivor pool is ∼70% (Fig. S3) which is consistent with serpentine tolerance being dominant to non-tolerance. Furthermore, the allele frequency in the non-serpentine survivor pool is ∼50% in this region as expected given that there were no survival differences at the non-serpentine site. The G-statistic and allele frequency estimates are somewhat noisy, largely due to the small size of the serpentine survivor pool which leads to increased sampling noise; this limited our ability to detect small effect QTL.

**Figure 3:**
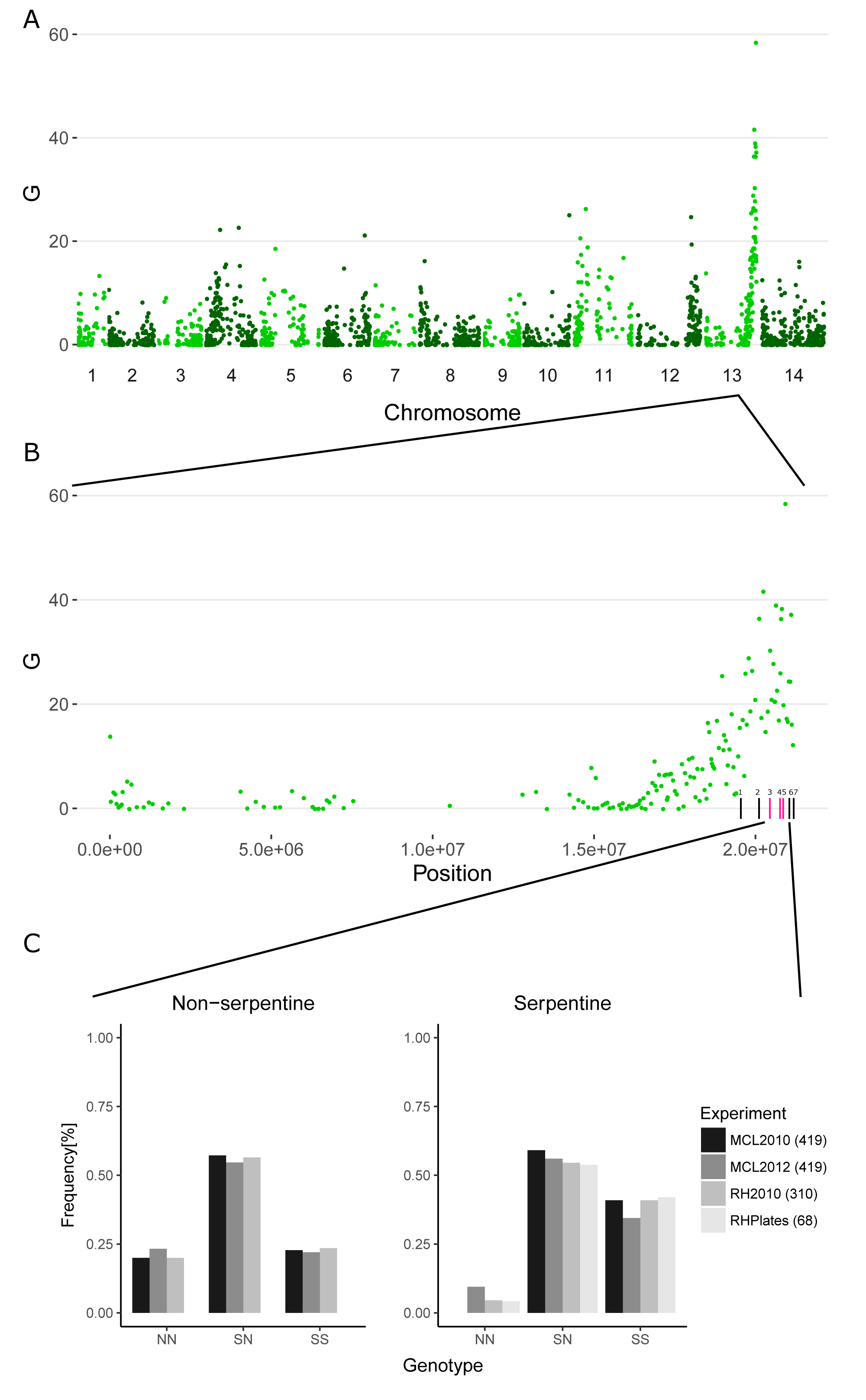
**A)** G-statistic calculated from summed allele counts in 25-SNP windows for the MCL2010 serpentine and non-serpentine survivor pools. Points indicate mean window position. **B)** G-statistic for chromosome 13 containing the highest peak. PCR-based markers used to genotype individual survivors for QTL confirmation shown on the X-axis. Markers from left to right are 778, 117, 310, 68, 419, 601 and 557 (MgSTS marker #′s http://www.mimulusevolution.org/). **C)** Genotype frequencies of survivors from field andcommon garden experiments shown for non-serpentine and serpentine sites. Survivors on serpentine soil deviate significantly from Mendelian expectations (1:2:1) in all experiments (MCL2010 (44): *χ*^*2*^*=*16.18, p=0.00031; MCL2012(116): *χ*^*2*^=16.19, p=0.00031; RH2010 (22): *χ*^*2*^=6.12,p=0.047; RHPlates (119): *χ*^*2*^=34.72, p=2.879e-08). The survivors at the non-serpentine field sites do not differ from expected (MCL2010 (215): *χ*^*2*^*=*4.8, p=0.091; MCL2012(236): *χ*^*2*^*=*2.13, p=0.35; RH2010 (85): *χ*^*2*^*=*1.635, p=0.442). Degrees of freedom=2 for all tests. Number of individual genotyped given in parentheses above. The marker ID used for each experiment provided in legend.

Using PCR-based markers (see Fig. 3B for marker locations) we genotyped all the individual survivors from both the serpentine and non-serpentine field sites for the MCL2010 experiment. The genotyping results confirm the putative QTL on chromosome 13 and show that it has a major effect on survival. As there were no significant differences in survival at the non-serpentine site we expect Mendelian segregation (1:2:1) in the survivors; however, survivors from the serpentine site will show segregation distortion at QTL controlling survival. Indeed, at one marker (MgSTS419) within the chromosome 13 QTL peak the non-serpentine survivors do not deviate from the expected 1:2:1 while none of the survivors at the serpentine site are homozygous for the non-serpentine allele (Fig. 3C). Other markers screened within the QTL region show similar patterns (sTable 4). Furthermore, the overall frequency of the serpentine allele was 69.8% in the serpentine survivors and their genotypic ratio was not significantly different from 1:2 serpentine homozygotes to heterozygotes (χ^2^=0.74, 1d.f., p=0.389), consistent with the serpentine allele being dominant. F3 survivors from the MCL2012 experiment were genotyped at the same maker and the serpentine survivors show significant distortion (Fig. 3C) while the non-serpentine survivors do not. Finally, we also show that this locus contributes to survival differences in the RH population where only a single survivor from the serpentine field sites is homozygous for the non-serpentine allele (Fig. 3C).

### QTL does not contribute to size differences at non-serpentine sites

The reciprocal transplant experiments found that non-serpentine plants were larger than serpentine plants in the non-serpentine field sites. To see whether the survival QTL on chromosome 13 contributes to these differences in plant size we conducted two-way ANOVAs in *lme4* for each of the three field experiments treating genotype as a fixed effect and block as a random effect. Only rosette diameter in the RH2010 experiment indicated a significant effect of genotype (sTable 5). A post-hoc Tukey test showed that the non-serpentine homozygotes were significantly smaller (0.76cm ± 0.09) than the heterozygotes (0.98cm ± 0.05) at p < 0.05. These size differences are in the opposite direction from what we observed in the field populations where plants from non-serpentine populations were larger. We do not know whether these juvenile size differences have an effect on fitness so it is not clear whether these genotypic differences are adaptive or maladaptive at the non-serpentine sites.

### Serpentine soils impose very strong selection

We calculated selection coefficients from the survivor genotype frequencies to understand how selection is acting on our QTL. Assuming initial genotype frequencies in the F2s/F3s were 1:2:1 at the time of transplanting, survival rates for each genotype were calculated by extrapolating the genotypic ratio from the survivors we were able to collect tissue from to the entire survivor pool. Using survival rate as our fitness measure, selection coefficients were calculated as 1-w12 or w22 for the relative fitness of the heterozygote and the non-serpentine homozygote respectively. Selection against the non-serpentine homozygotes was extremely strong in all three field experiments (Table 1) though it was weaker in the MCL2012 experiment likely due to the fact that this experiment was terminated 5 weeks earlier than the 2010 experiments. Serpentine tolerance is not completely dominant as the heterozygotes have a slightly lower survival rate compared to the serpentine homozygotes (Table 1) with dominance being less pronounced in the RH cross. These results demonstrate large and consistent effects of this QTL on serpentine survival across years and in different populations.

### Lab-based common garden replicates survival differences observed in field

Similar to the results from the field studies, the non-serpentine lines did not survive on the serpentine soils in the common garden experiment (Fig. 4). These results indicate that properties of the soils, as opposed to other environmental variables that differed between the field sites, are the primary selective agents contributing to the observed survival differences. Both serpentine lines had high survival on their home soils (Fig. 4). The MCL F1s and F2s had high survival on the MCL soil again indicating that tolerance is dominant in this cross. Furthermore, the MCL F2s 5 week survival rate was 83.6% consistent with the simple genetic basis of tolerance found by the field mapping study. The RH F2s had an intermediate survival rate on their home soil (40%). We genotyped these RH F2 survivors at a marker within our chromosome 13 QTL region and find that the genotype frequencies are significantly distorted relative to Mendelian expectations (Fig. 3C) and that the strength of selection is similar to that observed in the field (Table 1). Tolerance appears to be more partially dominant in the RH cross compared to MCL (Table 1) which may explain the overall lower survival rate of the RH F2s in the common garden set-up. Additionally, there may be other loci that contribute to survival on the RH soil.

**Figure 4:**
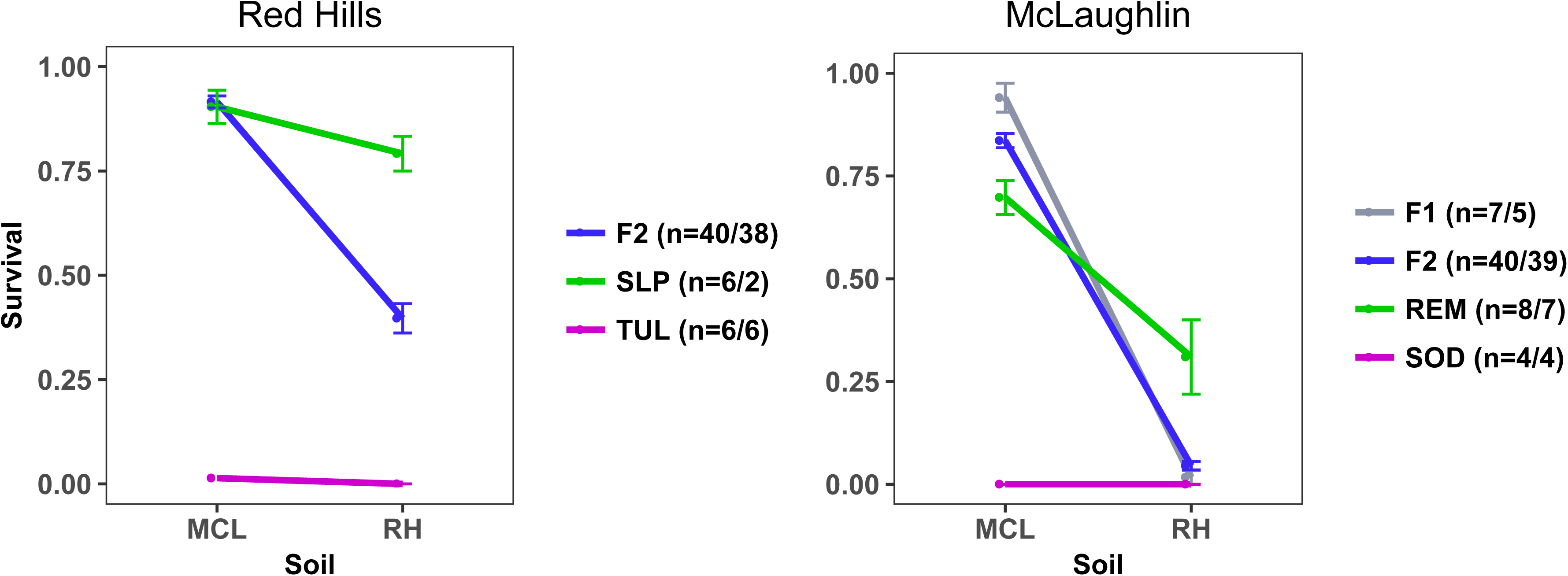
Survival in common garden experiment by genotype averaged across plates with standard errors. Serpentine populations shown in green and non-serpentine in pink. The number of plates planted per genotype on each soil type (MCL/RH soil) is given; 12 germinants planted/plate. The plot titles refer to the home region of the lines planted.

### Serpentine populations not equally tolerant of foreign serpentine soil

Despite the fact that the two serpentine lines share the major survival QTL they are not equally tolerant of each other′s soil. The RH serpentine line performs equally well on both soils (G=0.84, p=0.36) while the REM line has significantly lower survival on the RH soil than on its home soil (G=27.8, p=1.37e-7). Given that the seedlings were replicates of an inbred line this suggests that environmental variation contributes to survival. The differences between replicates could be due to variation in water availability both within and between plates, which would affect the rate at which plants are acquiring water and solutes from the soil matrix as well as heterogeneity in the soil itself. The non-serpentine lines had low survival on both soil types; however, the TUL line took longer to die on the MCL soil compared to the RH soil 100% mortality by week 1 census on RH soil but only 100% mortality at week 4 on MCL soil. The RH F2s had very high survival (94.1%) on the MCL soil which is likely due to this soil being less stressful to the non-serpentine parent TUL. The MCL F2s on the other hand had very low survival on the RH soil (4.5%). The difference in overall survival between the REM parental line and the MCL F2s on the RH soil indicates that there are other loci which contribute to survival on the RH soil. In addition, the REM and SLP lines may have different serpentine alleles at the QTL with different functionalities.

Variation in the chemical composition of the MCL and RH soils may help to explain the patterns of survival observed in the common garden experiment. The soil from the RH field sites has lower mean absolute levels of both Ca (112.83 ± 20.1) and Mg (1125 ± 34.5) as well as an overall lower Ca:Mg (0.1) compared to the MCL soils (Ca 515.17 ± 119.0; Mg 2471.83 ± 322.0; Ca:Mg 0.21). The RH soils also have higher average Ni ppm (19.5 ± 2.4) than MCL (8.4 ± 1.19) while the non-serpentine soils have low Ni levels (RH-off = 1.27 ±0.12; MCL-off 0.4 ± 0.08). Taken together these results suggest that the RH serpentine soil presents an overall harsher environment than the MCL soil. Elucidating the mechanism whereby soil chemical differences actually influence serpentine tolerance requires more detailed experiments isolating individual variables. Furthermore, a full QTL mapping study of the RH population would help to confirm whether there are other loci contributing to serpentine tolerance in this population.

## DISCUSSION

### Simple genetic basis of serpentine adaptation in M. guttatus

Non-serpentine populations of *M. guttatus* are unable to survive when planted on serpentine soils in both the field and the lab while serpentine populations enjoy moderate to high survival rates. Reciprocal transplant studies between serpentine and non-serpentine populations in other species have also found survival differences at serpentine sites (e.g. *Collinsia sparsiflora* (J. W. Wright *et al.*, 2006), *Helianthus exilis* (Sambatti & Rice, 2006), *Leptosiphon parviflorus* (Dittmar, 2017) and even long-lived pines (J. W. Wright, 2007)). We also found strong selection against non-serpentine plants in the common-garden experiment indicating that soil variables mediate these survival differences. By planting mapping populations in the field we were able to directly map loci contributing to these viability differences. The bulk segregant analysis identified a region on the end of chromosome 13 showing a large enrichment of serpentine alleles in the survivors from serpentine sites relative to survivors at the non-serpentine sites. Genotyping the individual F2 survivors from each of the habitats confirmed this region as a major effect QTL that explains 71% (2012) and 84% (2010) of the survival differences between the parents at the McLaughlin field sites. In addition, the serpentine allele is largely dominant with heterozygotes having only slightly reduced survival rates relative to serpentine homozygotes.

The simple genetic basis of serpentine adaptation in *M. guttatus* supports other QTL studies showing major gene effects for serpentine tolerance (Bratteler *et al.*, 2006; Burrell *et al.*, 2012). These QTL results contrast with findings from genome scans in *Arabidopsis* where many loci show elevated levels of differentiation between serpentine and non-serpentine populations (Arnold *et al.*, 2016; Turner *et al.*, 2010). However, it is necessary to connect variants to fitness differences in the field in order to understand their true adaptive value and by directly mapping on field survival differences, this study demonstrates that a major locus underlies adaptation to complex serpentine habitats in *M. guttatus*. We do not presume that this study presents the full picture of the genetic basis of serpentine adaptation in *M. guttatus* as our BSA was underpowered to detect loci of smaller effect due to the limited size of the serpentine survivor pool (Magwene *et al.*, 2011). Future work combining both high-powered mapping studies as well as genomic scan approaches will provide a more complete picture of the genetic architecture of adaption to serpentine soils in *M. guttatus*.

The QTL is currently localized to a roughly 1.5Mb region on the end of chromosome 13 which contains several hundred genes. This region contains a homolog of one of the putative serpentine adaptation genes in *A. lyrata* (Turner et al. 2010) that is in the RING/U-Box superfamily and is involved in zinc ion binding. A gene encoding a glutathione S-transferase (GST) protein which function in stress response and heavy metal tolerance is also found in this interval (reviewed Edwards *et al.*, 2000; Yadav, 2010). Additionally, a number of genes have annotations indicating transporter or metal binding activity. However, in order to prioritize candidate genes it will be necessary to identify the actual traits underlying the observed survival differences. Palm *et al.* (2012) grew the same REM and SOD lines as we used (these were the parents of the MCL mapping population) in hydroponic culture with altered Ca:Mg. They found that the serpentine line (REM) was more tolerant of the low Ca:Mg growth environment based on differences in biomass and phyotosynthetic rate. Adaptation to low Ca:Mg may be an important driver of adaptation to serpentine habitats in *M. guttatus.* However, our QTL interval does not contain any calcium or magnesium specific transporters such as CAX genes which have been implicated in tolerance to low Ca:Mg (Bradshaw, 2005). Finemapping efforts are underway that will narrow the QTL region to identify the causal locus. Finemapping will also help to address whether this QTL is actually comprised of multiple linked loci as might be predicted if adaptation to serpentine soils occurred and is maintained in the face of gene flow (Yeaman & Otto, 2011; Yeaman & Whitlock, 2011).

### Cost to tolerance

Local adaptation is defined as a genotype by environment interaction where local genotypes have higher fitness than foreign ones. The survival differences at the serpentine sites indicate strong fitness reductions for non-serpentine plants in these habitats. While there were no survival differences between the ecotypes at the non-serpentine field sites, we did detect differences in plant size where non-serpentine plants were larger than serpentine plants. Palm et al (2012) found similar differences in biomass between the REM/SOD pair from the McLaughlin Reserve when grown in potting soil. Work in other species has also found that serpentine-tolerant plants do not grow as well as non-serpentine plants when grown together on non-serpentine soils (Jurjavcic *et al.*, 2002; Kruckeberg, 1954; Proctor & Woodell, 1975; Sambatti & Rice, 2006). It′s thought that these growth rate differences may lead to a reduction in competitive ability of serpentine tolerant plants in non-serpentine sites which typically have higher vegetative cover. However, the connection between growth rate differences and fitness is not clear and we were limited in our ability to detect such a tradeoff as we were not allowed to let the transplants flower and set seed in the field.

### Same QTL contributes to serpentine adaptation in second population

The reciprocal transplant experiments found differences in survival rates between serpentine populations collected from different localities. At both MCL and RH the home serpentine populations had the highest survival suggesting that not all serpentine *M. guttatus* populations are equally tolerant to all serpentine soils but rather that they may be locally adapted to the specific characteristics of their home patch. However, these survival differences in the field could have been due to other environmental variables that differed between localities. The lab-based common-garden experiments directly tested the role of soil variables on survival. While the RH serpentine population enjoyed high survival on both soils, the MCL population had significantly lower survival when planted on the RH soil compared to its home soil. Such findings are perhaps not surprising given the patchy distribution of serpentine soils and variation in chemical (Fig. 1) and physical properties arising from differences in the primary mineralogical composition of parent materials, degree and conditions of metamorphic alteration and degree of weathering (Alexander *et al.*, 2007; Kruckeberg, 1984; Proctor *et al.*, 1975; Whittaker, 1954).

The tolerance differences between the RH and MCL populations are especially interesting because we found that they share the same major QTL. Genotyping the survivors from the RH serpentine field sites and soil plates showed that very few of the survivors were homozygous for the non-serpentine allele at the QTL (s=0.9 in both experiments). We do not know whether the RH and MCL populations represent independent evolutions of serpentine tolerance. However, based on geography – these two populations are ∼300km apart and separated by the Central Valley in CA where there is no serpentine soil (Fig. 1) – this seems like a reasonable hypothesis. The differences in survival on the two soils between the populations suggests that either there are other loci required to grow on the RH soil which are not shared between the populations or that the populations have different alleles, or even different loci underlying this shared QTL. Serpentine and non-serpentine populations of *M. guttatus* grow throughout Western North America, often in close proximity, providing an ideal system for investigating the distribution of this major serpentine tolerance QTL and to address questions of parallel adaptation. Future work aims to determine whether the shared genetic basis of tolerance is due to a single mutational origin followed by migration or if there has been repeated selection from standing variation or independent mutations at the same locus.

## Conclusion

Many classic examples of adaptive differentiation between populations or closely related species involve changes at one or a few loci: cryptic coloration (Hoekstra *et al.*, 2006; van′t Hof *et al.*, 2011), mimicry (Baxter *et al.*, 2010), shifts from marine to freshwater (Colosimo *et al.*, 2005) and pollinator shifts (Bradshaw Jr & Schemske, 2003). These cases are similar to the colonization of serpentine soils where populations may initially be far from a new fitness optima with limited intermediate habitat and/or where intermediate phenotypes have low fitness. Such situations likely favor the fixation of large effect loci (Dittmar *et al.*, 2016). *M. guttatus* has colonized other harsh habitats such as copper mine tailings, hot springs and coastal cliffs (Selby *et al.*, 2014) and large effect QTLs have been found for traits contributing to adaptation to these habitats (Hendrick *et al.*, 2016; Lowry *et al.*, 2009; K. M. Wright *et al.*, 2015).

Interestingly, none of these major QTL involved in adaptation to stressful abiotic habitats in *M. guttatus* are shared. For example, the major copper tolerance locus is on chromosome 9, yet, the focal copper tolerant populations occur only 20km from the RH serpentine site. While we have not investigated whether there may be some degree of cross tolerance between serpentine and copper adapted populations, they appear to have largely colonized these harsh habitats via independent, large effect loci. How *M. guttatus* consistently has the necessary genetic variation to produce these large phenotypic shifts to colonize numerous harsh habitats while many other species occurring in close proximity fail to adapt is unknown.

## Data Accessibility

All data will be made publicly available on Dryad upon the acceptance of this manuscript.

## Author Contributions

JPS and JHW designed the experiments and wrote the paper. JPS performed all experiments and analyses.

## Acknowledgements

Thank you to members of the Willis lab past and present and Graham Coop for comments on the ideas presented within and drafts of this manuscript. In particular, thank you to Kathy Toll for field assistance in 2012. We are also extremely grateful to Cathy Koehler and Paul Aigner at the McLaughlin Reserve for their help facilitating the field experiments and providing housing. We would also like to thank Sonya Jooma who assisted with plant care, field work and genotyping and Lex Flagel and Chenling Xu for help with the initial analysis of the bulk segregant data. This work was supported by grant number NSF IOS-1354688 to JHW.

